# Dosing regimen of gentamicin and the effect of antenatal steroids on clearance in preterm infants

**DOI:** 10.1101/566125

**Authors:** Roberto P. Santos, Yin Chang, Carlos R. Oliveira, Beverly Adams-Huet, John Gard, Pablo J. Sanchez

## Abstract

**Objective:** Determine the gentamicin pharmacokinetic and pharmacodynamic (PK/PD) profiles of once daily dosing (ODD) versus every 18-hour (q18h) regimens and to characterize the effect of antenatal steroid on gentamicin clearance (Gent**_CL_**).

**Study Design:** Retrospective cohort of preterm infants (≤34 weeks gestation; ≤7 days) who received gentamicin for >48 hours from January 2005 to June 2007. Serum gentamicin peak and trough concentrations were determined, and PK/PD profiles calculated using standard noncompartmental methods.

**Result:** 122 (63%) infants received ODD and 73 (37%) received q18h regimen. Desired gentamicin peak (5 to 12 mcg/mL) and trough (<2 mcg/mL) concentrations were achieved in 80% (95%CI, 72-86) on ODD vs. 47% (95%CI, 36-58) on q18h (*p<0.001*). Target drug exposure (AUC >72 mcg/mL/hr) was achieved in 73% (89/122) of infants on ODD vs. 22% (16/73) on q18h (p < 0.001). Gent**_CL_** was significantly lower in those who receive antenatal steroid (37+/-8 mL/kg/hr vs. 42+/-13 mL/kg/hr, *p=0.04)* but not affected by postnatal indomethacin treatment (*p>0.86*).

**Conclusion:** PK/PD profile of gentamicin is improved by ODD in preterm infants. Gent**_Cl_** was significantly less in infants exposed to antenatal steroids but not in those treated with indomethacin.

## Introduction

Gentamicin is the preferred antimicrobial agent for the treatment of Gram-negative infections in neonates.^1,2^ Therapeutic drug monitoring is needed since it has a narrow therapeutic index.^3-5^ Once daily dosing (ODD) of gentamicin (4 mg/kg IV q24 hours) has been used in the normal newborn nursery at Parkland Memorial Hospital (PMH) since 2000.^6-8^ The pharmacodynamic characteristics of aminoglycosides that allow the use of once-daily dosing include concentration dependent (C**_max_**/MIC ratio) killing,^2,3,9^ post-antibiotic effect with leukocyte enhancement,^10,11^ and prevention of adaptive resistance.^10,12,13^

Some institutions still utilize other extended interval dosing regimen for gentamicin among preterm infants. For 30-34 weeks gestation ≤7days old, gentamicin is given at 4.5 mg/kg every 36 hours (hrs) and for ≤29 weeks gestation ≤7days old 5 mg/kg every 48 hrs is recommended.^14,15^ Since December 2005, PMH had changed to ODD dosing regimen of gentamicin for preterm infants 27-34 weeks gestation at 3 mg/kg and for ≤26 weeks gestation at 2.5 mg/kg with good pharmacokinetic-pharmacodynamic profiles.^7^

Gentamicin clearance (Gent**_CL_**) correlates with glomerular filtration rate (GFR),^2,3,16^ which in turn may be affected by prenatal exposure to maternal corticosteroids given for promotion of lung maturation. Antenatal exposure to dexamethasone may enhance renal tubular sodium reabsorption immediately after birth.^17^ This may be mediated by earlier maturation of renal tubules through activation of tubular Na+-K+ ATPase enzymatic system.^17^

Prenatal exposure to indomethacin or betamethasone can significantly affect the renal function of preterm infants after birth. Antenatal exposure to betamethasone with indomethacin may significantly increase GFR.^18^ Previous reports in rat model suggest that glucocorticoid-induced (methylprednisolone) increase GFR is associated with increase in glomerular plasma flow rate.^19,20^ However, the effect of prenatal exposure to indomethacin on GFR or on Gent**_CL_** when given simultaneously with steroid has not been fully elucidated.

The main objective of this study was to determine the gentamicin pharmacokinetic (PK)-pharmacodynamic (PD) profiles of ODD versus q18h dosing regimens and to characterize the effect of antenatal steroid exposure on Gent**_CL_**. We hypothesized that ODD of gentamicin has better PK-PD profiles than q18h regimen, and that antenatal exposure to steroids affects Gent**_CL_** (a surrogate marker of GFR).

## Methods

This is a retrospective cohort study of preterm infants gestational age ≤34 weeks gestation and ≤7 days old who received gentamicin intravenously for >48 hrs for possible Gram-negative infection. This was done at the normal newborn nursery and nursery intensive care unit of PMH from January 2005 – June 2007. During those periods, PMH has an estimated 16,000 deliveries annually. Medical records were reviewed and abstracted for demographic and clinical data.

On December 2005, administration of gentamicin to preterm infants (gestational age 27-34 weeks) in the neonatal intensive care unit at PMH changed from 2.5 mg/kg q18h dosing regimen to ODD (3 mg/kg). Infants with GA ≤ 26 wk continued to receive ODD (2.5 mg/kg). Serum creatinine (S_crea_) was performed as part of the patient’s medical care and creatinine clearance (Crea_CL_) was estimated using Schwartz method.^21^

Gentamicin peak (1 hr) and trough (23 hrs) concentrations were obtained after the 3^rd^ dose considered the steady state per PMH nursery protocol. The serum peak and trough concentrations were determined by fluorescence polarization immunoassay.^3^ The sensitivity limit of this assay (AxSYM, Abbott Laboratories, Abbott Park, IL, USA) is 0.3 mg/L with coefficient of variation <5%.^15^

Pharmacokinetic profiles were determined using standard non-compartmental methods.^22,23^ (Appendix 1) The target PD indices such as area under the concentration-time curve (drug exposure, AUC)^3^ and the maximum concentration/MIC ratio, (C**_max_**/MIC)^2,3,9,14^ were calculated.

Maternal steroid regimen administered antenatally from January 2005 to December 2005 was dexamethasone 5 mg every 12 hours (4 doses) and from January 2006 to June 2007 it was betamethasone 12 mg every 24 hours (2 doses). Indomethacin was given to infants ≤4 days old as standard of care for medical management of patent ductus arteriosus. The dose was 0.2 mg/kg intravenously every 12 hours (3 doses).

No infant had a positive blood or cerebrospinal fluid culture for Gram negative organisms during the study period and thus, microbiologic efficacy was not determined.

Descriptive analyses (means, standard deviations, proportions), confidence intervals, and two-way ANOVA on ranks and multiple comparisons were used to determine the steroid effect on Gent**_CL_** and interaction analysis between steroid and indomethacin were done (SAS Institute Inc. v9.1, Cary, NC) with *p<0.05* (2 sided) considered significant.

This medical record review was approved by the Institutional Review Board of the University of Texas Southwestern Medical Center as part of the research project, “nomogram for gentamicin dosing in early infancy” (IRB File Number 012006-050).

## Results

195 preterm infants (103 male) gestational age (mean ± SD) 30 ± 3 weeks with mean birth weight, 1591 ± 601 grams were enrolled at 3 ± 1 days of age. 122 (63%) infants received ODD and 73 (37%) received q18h regimen. (Figure 1) 25 infants who received ODD gentamicin had a mean S_crea_, 1.0 ± 0.2 mg/dL and a mean Crea_CL_ of 12.6 ± 3.6 mL/min per 1.73 m^2^. 20 infants who received q18h gentamicin had a mean S_crea_, 1.2 ± 0.1 mg/dL and a mean Crea_CL_ of 10.8 ± 0.2 mL/min per 1.73 m^2^.

**Figure 1.**
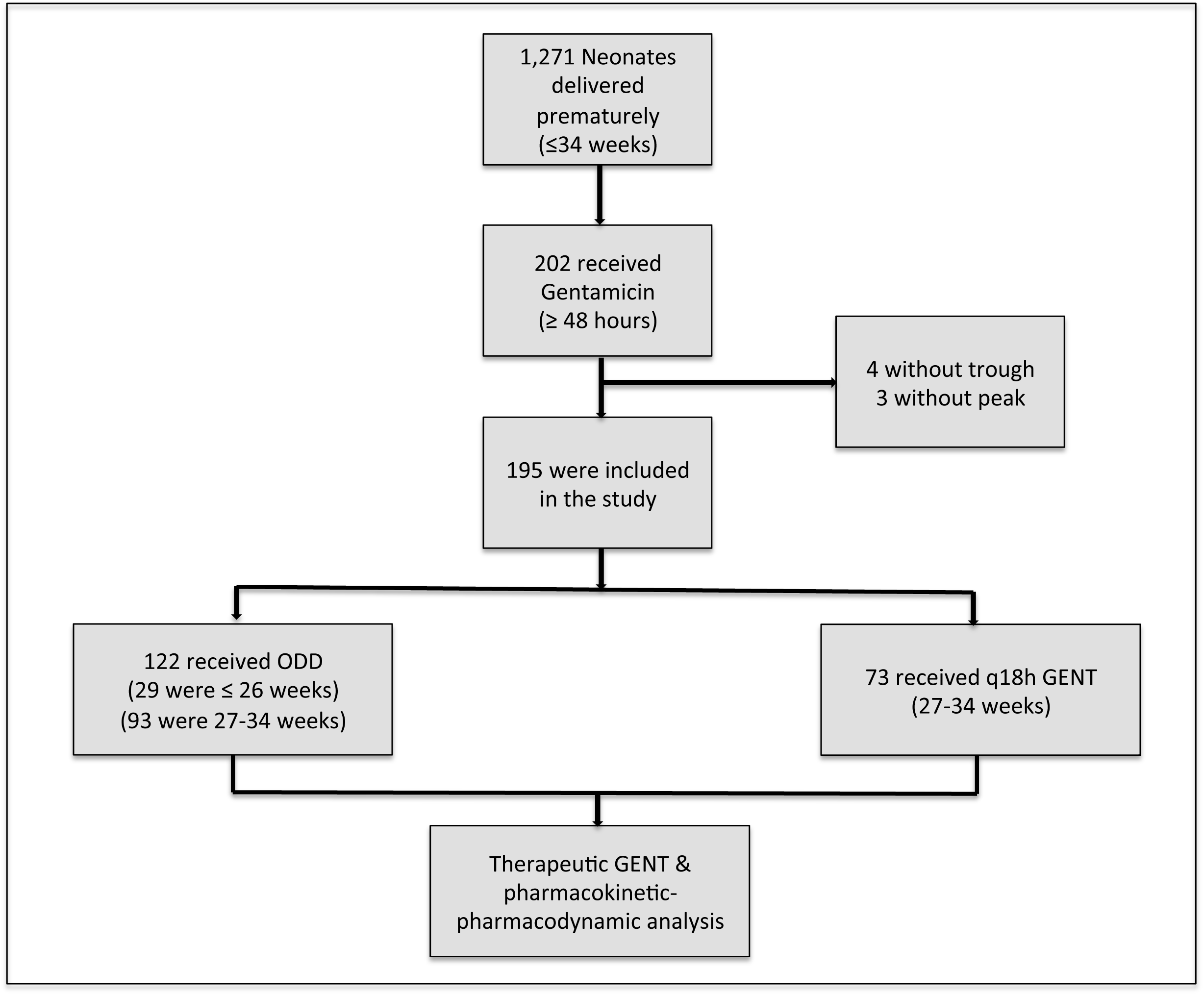
Enrollment for therapeu0c gentamicin (GENT) and pharmacokine0c;pharmacodynamic profile of once daily dosing (ODD) versus every 18 hour (q18h) regimen

### A. PK-PD profiles of gentamicin ODD versus every 18 hours

Desired gentamicin peak (5-12 mcg/mL) and trough (<2 mcg/mL) concentrations^2,3,14,15^ were achieved in 80% (95%CI, 72-86) on ODD vs. 47% (95%CI, 36-58) on q18h (*p<0.001*). (Figure 2)

**Figure 2.**
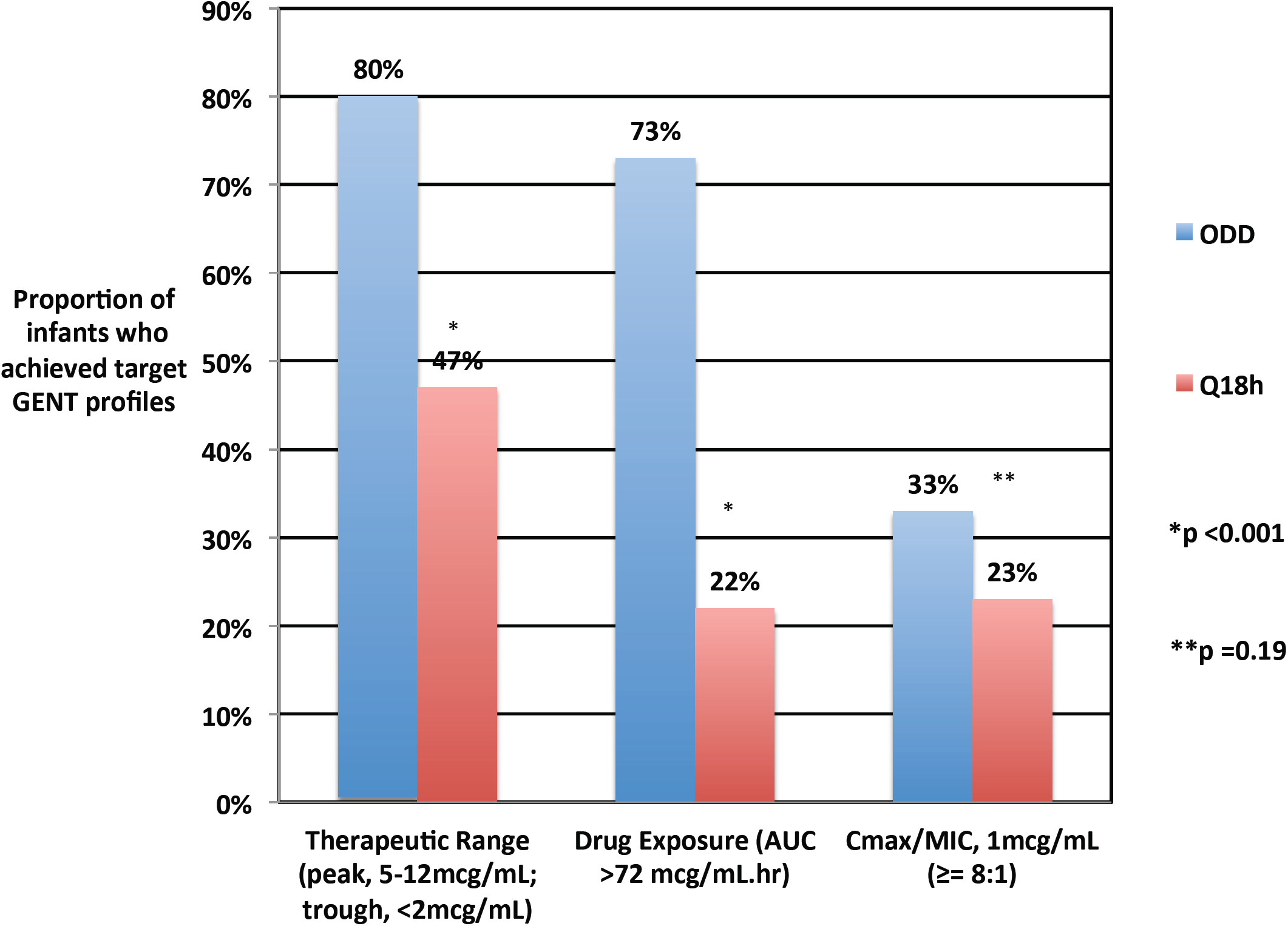
Gentamicin (GENT) pharmacokinetic<pharmacodynamic profiles for once daily dosing (ODD) versus every 18 hour (q18h) regimen in preterm infants.

The PK profile of ODD includes Ke 0.07 ± 0.02 h**^- 1^**, t**_1/2_** 10.3 ± 2.4 hrs, C**_max_** 7 ± 1.4 mcg/mL, C**_min_** 1.4 ± 0.5 mcg/mL, Vd 0.5 ± 0.2 L/kg, CL 38.6 ± 11.5 mL/kg·hr, and AUC 82 ± 8 mcg·mL/hr. The PK profile of q18h showed Ke 0.08 ± 0.03 h**^- 1^,** t**_1/2_** 9.1 ± 2.9 hrs, C**_max_** 6.7 ± 1.2 mcg/mL, C**_min_** 1.7 ± 0.6 mcg/mL, Vd 0.5 ± 0.2 L/kg, CL 42.2 ± 10.4 mL/kg·hr, and AUC 63 ± 13 mcg·mL/hr.

Target drug exposure (AUC 72 mcg/mL hr)^3^ was achieved in 73% (89/122) of infants on ODD vs. 22% (16/73) of those on the q18h regimen (*p < 0.001).* (Figure 2)

A desired C**_max_**/MIC (1 mcg/mL) ratio of 8:1^2,3,9,14^ was achieved in 33% (95%CI, 25-42) of infants on ODD vs. 23% (95%CI, 15-34) on q18h regimen (*p = 0.19*). (Figure 2)

### B. PK-PD profiles of gentamicin ODD: Gestational age ≤26 weeks versus 27-34 weeks

Desired gentamicin peak (5-12 mcg/mL) and trough (<2 mcg/mL) concentrations^2,3,14,15^ (*p =0.06*), the target drug exposure (AUC 72 mcg/mL hr)^3^ (*p =0.17*), and the desired C**_max_**/MIC (1 mcg/mL) ratio of 8:1^2,3,9,14^ (*p =0.23*) were not significantly different between gestational age ≤26 weeks versus 27-34 weeks using ODD gentamicin. (Figure 3)

**Figure 3.**
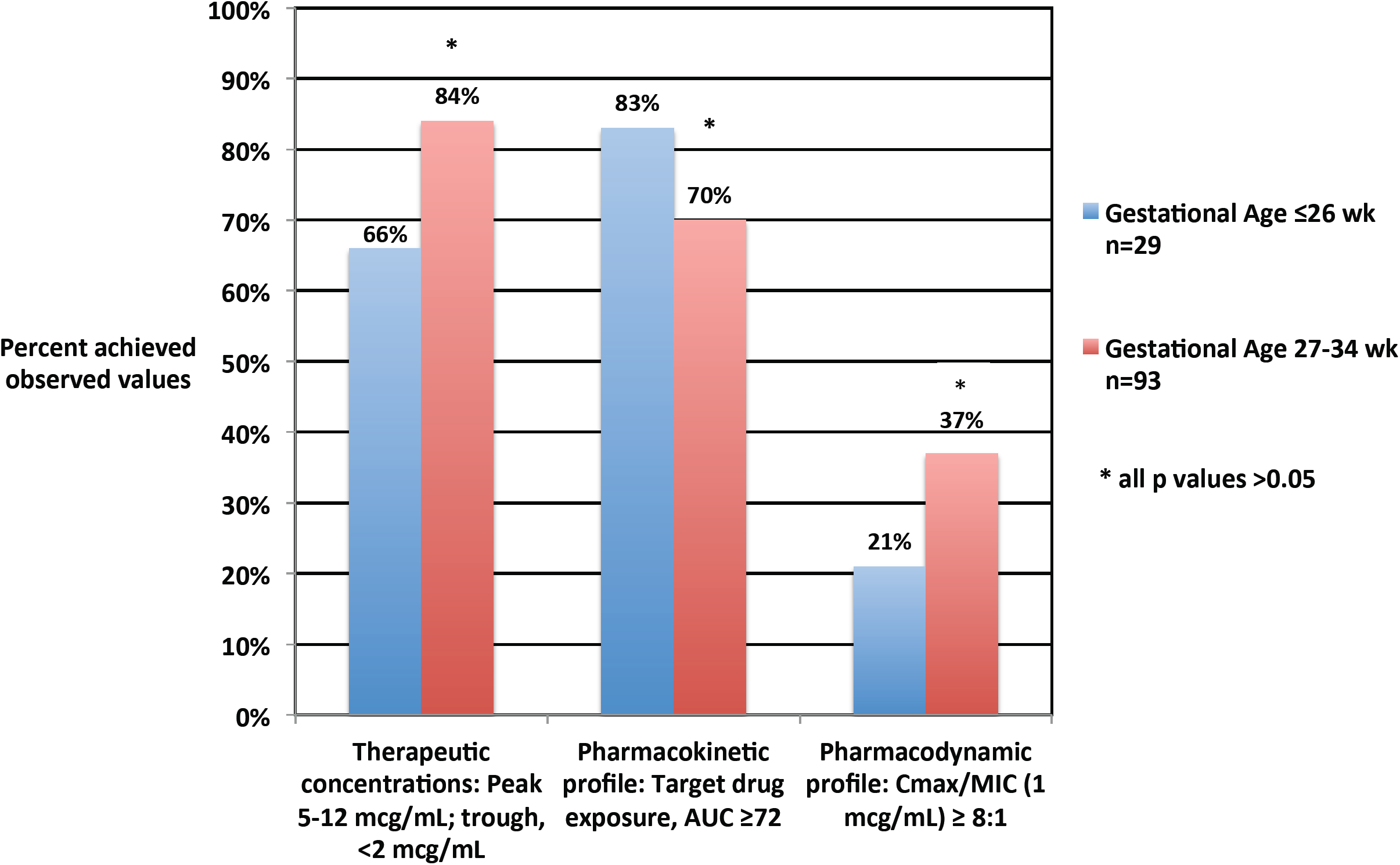
Gentamicin pharmacokinetic-pharmacodynamic profiles for once daily dosing regimen for gestational age ≤26 weeks versus 27=34 weeks.

### C. Gentamicin clearance according to antenatal exposure to steroid with or without postnatal exposure to indomethacin

129 (66%) infants did not receive either steroid or indomethacin (Group A), 42 (22%) was exposed to steroid antenatally only (Group B), 14 (7%) received indomethacin postnatally (Group C), and 10 (5%) was exposed to steroid antenatally and received indomethacin postnatally (Group D). (Figure 4)

**Figure 4.**
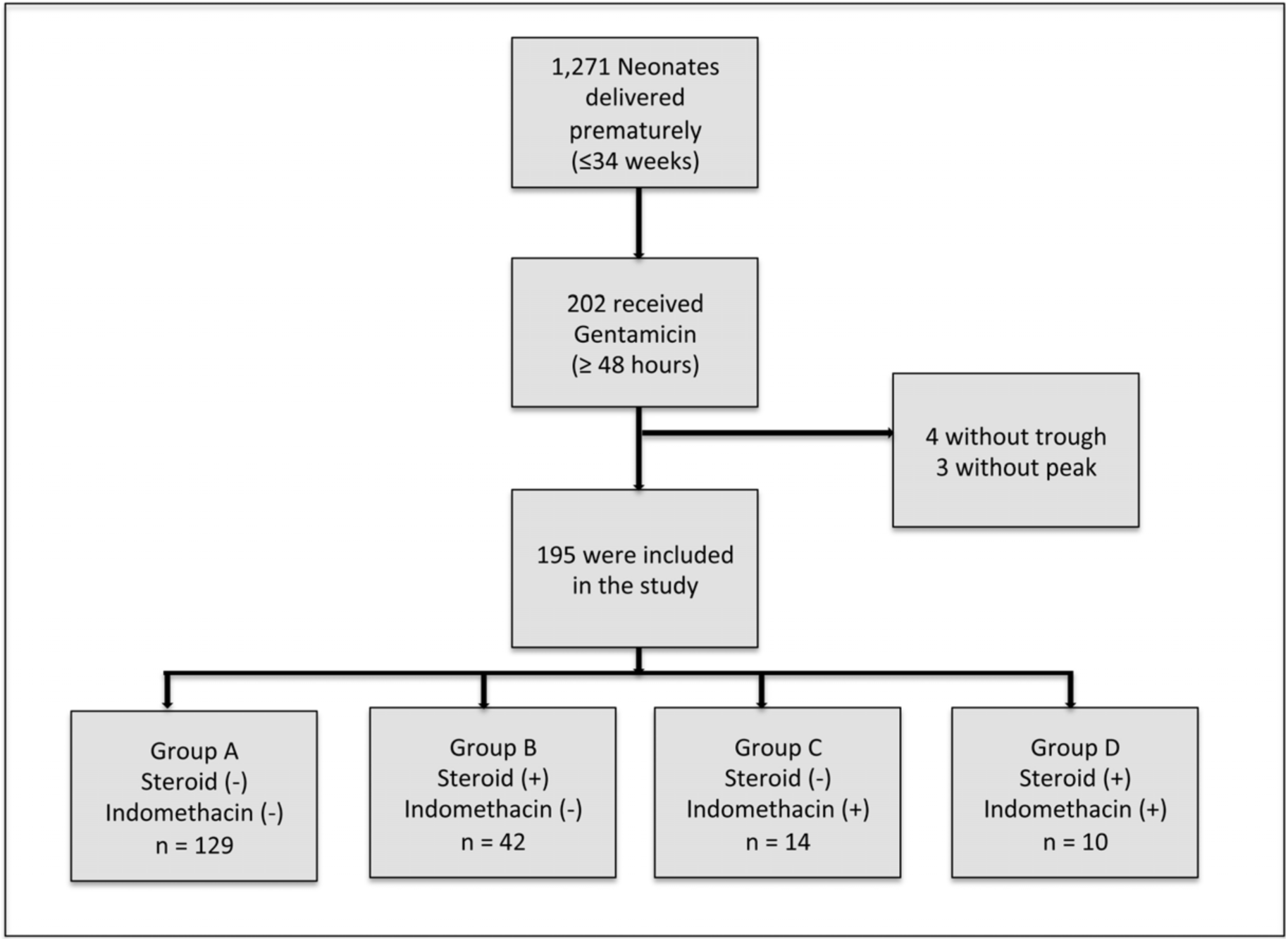
Enrollment for gentamicin clearance according to prenatal exposure to steroid, postnatal exposure to indomethacin, or both

There was a trend for decrease Gent**_CL_** in preterm infants that received antenatal steroid alone (Group B, Gent**_CL_** = 37.1 ± 8.2, p=0.06) and those who received antenatal steroid and postnatal indomethacin (Group D, Gent**_CL_** = 36.4 ± 7.1, p=0.28) compared to those who did not receive either medications (Group A, Gent**_CL_** = 41.3 ± 12.5).

Gent**_CL_** was significantly lower among infants exposed to antenatal steroids than in those whose mothers did not receive steroids (37 ± 8 mL/kg·hr vs. 42 ± 13 mL/kg·hr, *p = 0.04).* The mean decrease in Gent**_CL_** was approximately 4.6 ± 8 mL/kg·hr. There was no significant steroid and indomethacin interaction (*p>0.86*).

### D. Clinical response and safety

All patients improved clinically in response to treatment with ampicillin and gentamicin for neonatal sepsis regimen. No patients had Gram negative bacteremia, meningitis, or brain abscess.

All patients tolerated gentamicin and there were no severe adverse events requiring drug discontinuation. Of those patients with available serum creatinine values as part of their medical care while on gentamicin, only one developed renal failure that eventually died. Majority of patients passed their newborn hearing evaluations except for three infants who failed the initial hearing test and on follow up one was diagnosed with retrocochlear pathology, another showed normal hearing evaluation, and in another the repeat hearing test was not performed because of death.

Microbiologic efficacy cannot be determined since no Gram negative organism was isolated during the study period thus, actual minimum inhibitory concentration cannot be used for PD calculations.

## Discussion

The pharmacokinetic and pharmacodynamic profiles of gentamicin are improved by ODD in preterm infants. ODD gentamicin regimen achieved desired therapeutic concentrations and PK-PD profiles compared to every 18 hour dosing regimen in preterm neonates. Gentamicin clearance was significantly affected in infants exposed to antenatal steroids but not in those postnatally treated with indomethacin.

The desired PK-PD profiles were better using ODD of gentamicin in our patients compared to q18 hr regimen. More patients achieved the desired peak and trough using ODD regimen. Because of the narrow therapeutic index of gentamicin^3-5^ we continue to follow the gentamicin peak and trough concentrations. These values are readily available in most clinical settings with results available usually after several hours. These therapeutic ranges may be followed by clinicians to guide their clinical management and individualize treatment.^3^

There were more preterm infants who achieved the target drug exposure using ODD of gentamicin. It was previously suggested that gentamicin AUC approach should be followed in dose-individualization.^3^ This was based on the principle that the same total dose given over 24 hrs should be given using individualization of therapy with the conventional or multiple dosing regimens. Assuming a one-compartment model, drug exposure can be calculated for each patient and subsequent doses may be adjusted to achieve target AUC which is associated with the efficacy of aminoglycosides.^2^

We used a desired C**_max_**/MIC ratio of 8:1 since this was recommended as the useful range of bactericidal activity against most bacteria. This bactericidal effect is usually about 5-10 times the MIC of the organism.^2,3,9,14^ In our patients, there was a trend to achieve this PD index using the ODD regimen compared to q18h.

The desired gentamicin therapeutic ranges (peak and trough), target drug exposure and a desired C**_max_**/MIC ratio were very similar between preterm newborns ≤26 weeks and those infants 27-34 weeks gestation. This strongly suggests that desired PK-PD indices may be achieved using ODD regimen in preterm infants. There was a trend to achieve more target drug exposure among infant ≤26 weeks gestation than those born at 27-34 weeks. This may be associated with longer half-life of gentamicin which may approach >10 hours among very low birth weight infants with gestational age <31 weeks.^15^ In preterm infants this may be due to larger extracellular fluid and volume of distribution resulting in reduced clearance.^2,16^ This may significantly affect the drug exposure (AUC = dose given/ Cl) among infants with very low birth weight.

There are extended interval dosing regimens other than ODD for gentamicin. Lanao, J.M., et al. had described the PK basis for the use of extended interval dosing regimen of gentamicin for neonates using population PK modeling to achieve desired serum concentrations.^15^ However, these findings should be correlated with clinical outcome and patient safety as well as microbiologic efficacy. One of the strengths of our research was to determine not only the clinical outcome but the patient safety as well with ODD regimen through a retrospective cohort design. We were not able to establish microbiologic efficacy since no infant had a positive blood or cerebrospinal fluid culture for Gram negative organisms. Thus, we were unable to calculate the PD profile using actual MIC values. However, we estimated the desired PD index of C**_max_**/MIC ratio of ≥ 8:1 ^2,3,9,14^ assuming an MIC value of 1 mcg/mL.

Gent**_CL_** was significantly affected by antenatal steroid exposure. In our patients, contrary to previous reports,^17,18^ exposure to dexamethasone or betamethasone significantly decreased Gent**_CL_**. The effect was consistent in the groups of infants that were exposed to antenatal steroid without postnatal indomethacin (Group B) and in those who were exposed to antenatal steroid with postnatal indomethacin (Group D). (Figure 4) There was no evidence to support that the exposure to steroid and indomethacin in decreasing Gent**_CL_** was synergistic or additive. Our finding does not support previous studies on enhanced Gent**_CL_** associated with antenatal exposure to steroid.^17,18^

The importance of this study has implications in guiding clinicians in the therapeutic management of preterm infants. ODD gentamicin may be more practical and convenient than every 36 hours or every 48 hours regimen particularly in preterm infants receiving other medications. The simplicity & practicability of ODD regimen may potentially avoid missing of doses and prevent medication errors. This however, requires prospective evaluation.

This may increase awareness among health providers that medications given antenatally may significantly affect Gent**_CL_** in preterm newborn postnatally. The effect of steroid given antenatally may increase clinical suspicion and be vigilant in monitoring renal insufficiency while being administered potentially nephrotoxic agents such as vancomycin, furosemide, and aminoglycosides.

The findings in this study are subject to several limitations. First, the inherent nature of a retrospective study may only suggest association and does not allow causality of the effect of antenatal steroid exposure on Gent**_CL_**. However, the sample size was large enough (N = 195) to detect a change of >4.5 mL/kg·hr in Gent**_CL_** (α=0.01, β<0.05).^24^ Second, there were two steroid regimens used during the study period (dexamethasone in January-December 2005 and bethamethasone thereafter starting in January 2006) which does not allow us to separate the effect of either one on Gent**_CL_**. Previous studies have reported prenatal exposure to dexamethasone^17^ or betamethasone^18^ may exert significant effect on the enhanced maturation of renal function in preterm infants. Both synthetic glucocorticoid have equivalent potency.^18^ Finally, we did not control or take into consideration other medications that the infants were receiving during the study period or clinical parameters that may affect the renal function such as sepsis, patent ductus arteriosus (PDA), and those on extracorporeal membrane oxygenation (ECMO).^2^ These conditions (severe infections, PDA, ECMO) have been associated with increase in volume of distribution requiring higher dosing regimen of gentamicin to achieve desired peak concentrations.^3^

In conclusion, gentamicin ODD regimen is clinically effective and safe among preterm infants (2.5 mg/kg for ≤26 weeks gestation and 3 mg/kg for 27-34 weeks). The PK-PD profiles (desired peak/ trough, target drug exposure, and desired C**_max_**/MIC ratio) were achieved among preterm infants with ODD. Antenatal steroid exposure can significantly affect Gent**_CL_** of preterm infants that cannot be explained by indomethacin exposure postnatally. We recommend prospective evaluation of the PK-PD profile of ODD gentamicin regimen in preterm infants and correlating them with clinical outcome and microbiologic efficacy, as well as the effect of antenatal steroid exposure on Gent**_CL_**.

## Acknowledgements

We acknowledged the technical and statistical support provided by the Department of Clinical Sciences (Master’s Program in Clinical Sciences), University of Texas Southwestern Medical Center.

## References

1. Pickering L, ed Red Book. 29th ed. Elk Grove Village: American Academy of Pediatrics; 2012. CJ Baker DK, SS Long ed. Escherichia coli and Other Gram-Negative Bacilli.

2. de Hoog M MJ, van den Anker JN. The use of aminoglycosides in newborn infants. In: I choonara AN, G Kearns, ed. Introduction to Paediatric and Perinatal Drug Therapy. Notthingham: Nottingham University Press; 2003:117–140.

3. Begg EJ, Barclay ML, Kirkpatrick CJ. The therapeutic monitoring of antimicrobial agents. Br J Clin Pharmacol. Jan 1999;47(1):23–30.

4. Fanos V, Dall’Agnola A. Antibiotics in neonatal infections: a review. Drugs. Sep 1999;58(3):405–427.

5. Paap CM, Nahata MC. Clinical pharmacokinetics of antibacterial drugs in neonates. Clin Pharmacokinet. Oct 1990;19(4):280–318.

6. Santos R GJ, Adams-Huet B, Ruparel J, Leff R, Bush M, Martin K, Tharayl S, Olesen B, Vedro D, Vinson J, Stehel E, Sanchez PJ. Validation of a random concentration-time curve (Gent-Curve) for once daily dosing of gentamicin among infants >/=35 weeks gestation. Pediatric Academic Societies & Asian Society for Pediatric Research; 2007; Toronto, Canada.

7. Santos R CY, de Oliveira C, Huet-Adams B, Gard J, Sanchez P. Dosing regimen of gentamicin and effect of antenatal steroids on clearance in preterm infants. Abstract presented at 2008 Pediatric Academic Societies’ & Asian Society for Pediatric Research Joint Meeting; May 2-6, 2008, 2008; Honolulu, Hawaii.

8. Jackson GL, Sendelbach DM, Stehel EK, Baum M, Manning MD, Engle WD. Association of hypocalcemia with a change in gentamicin administration in neonates. Pediatr Nephrol. Jul 2003;18(7):653–656.

9. Moore RD, Lietman PS, Smith CR. Clinical response to aminoglycoside therapy: importance of the ratio of peak concentration to minimal inhibitory concentration. J Infect Dis. Jan 1987;155(1):93–99.

10. Craig WA. Once-daily versus multiple-daily dosing of aminoglycosides. Journal of chemotherapy. Jun 1995;7 Suppl 2:47–52.

11. Novelli A, Mazzei T, Fallani S, Cassetta MI, Conti S. In vitro postantibiotic effect and postantibiotic leukocyte enhancement of tobramycin. Journal of chemotherapy. Aug 1995;7(4):355–362.

12. Rotschafer JC, Zabinski RA, Walker KJ. Pharmacodynamic factors of antibiotic efficacy. Pharmacotherapy. 1992;12(6 Pt 2):64S-70S.

13. Mattie H, Craig WA, Pechere JC. Determinants of efficacy and toxicity of aminoglycosides. J Antimicrob Chemother. Sep 1989;24(3):281–293.

14. PDR Network L, ed Neofax 2011. 24th ed. Montvale: Thomson Reuters; 2011. Gentamicin.

15. Lanao JM, Calvo MV, Mesa JA, et al. Pharmacokinetic basis for the use of extended interval dosage regimens of gentamicin in neonates. J Antimicrob Chemother. Jul 2004;54(1):193–198.

16. Koren G, James A, Perlman M. A simple method for the estimation of glomerular filtration rate by gentamicin pharmacokinetics during routine drug monitoring in the newborn. Clin Pharmacol Ther. Dec 1985;38(6):680–685.

17. Zanardo V, Giacobbo F, Zambon P, et al. Antenatal aminophylline and steroid exposure: effects on glomerular filtration rate and renal sodium excretion in preterm newborns. J Perinat Med. 1990;18(4):283–288.

18. van den Anker JN, Hop WC, de Groot R, et al. Effects of prenatal exposure to betamethasone and indomethacin on the glomerular filtration rate in the preterm infant. Pediatr Res. Nov 1994;36(5):578–581.

19. Baylis C, Brenner BM. Mechanism of the glucocorticoid-induced increase in glomerular filtration rate. Am J Physiol. Feb 1978;234(2):F166–170.

20. DeBermudez L, Hayslett JP. Effect of methylprednisolone on renal function and the zonal distribution of blood flow in the rat. Circ Res. Jul 1972;31(1):44–52.

21. Brion LP, Fleischman AR, McCarton C, Schwartz GJ. A simple estimate of glomerular filtration rate in low birth weight infants during the first year of life: noninvasive assessment of body composition and growth. The Journal of pediatrics. Oct 1986;109(4):698–707.

22. Atkinson AJ DC, Dedrick RL, Markey SP, ed Principles of Clinical Pharmacology. San Diego: Academic Press; 2007.

23. Winters M, ed Basic Clinical Pharmacokinetics. Philadelphia: Lippincott Williams & Wilkins; 2004.

24. Statistical power calculations online for two means. 1998.

